# Increased pericarp cell length underlies a major QTL for grain weight in hexaploid wheat

**DOI:** 10.1101/117937

**Authors:** Jemima Brinton, James Simmonds, Francesca Minter, Michelle Leverington-Waite, John Snape, Cristobal Uauy

**Affiliations:** John Innes Centre, Norwich Research Park, Norwich, NR4 7UH, United Kingdom

**Keywords:** wheat, thousand grain weight, yield, grain size, grain length, cell size, pericarp, QTL

## Abstract

- Crop yields must increase to address food insecurity. Grain weight, determined by grain length and width, is an important yield component, but our understanding of the underlying genes and mechanisms is limited.
- We used genetic mapping and near isogenic lines (NILs) to identify, validate and fine map a major quantitative trait loci (QTL) on wheat chromosome 5A associated with grain weight. Detailed phenotypic characterisation of developing and mature grains from the NILs was performed.
- We identified a stable and robust QTL associated with a 6.9 % increase in grain weight. The positive interval leads to 4.0 % longer grains, with differences first visible twelve days post fertilization. This grain length effect was fine-mapped to a 4.3 cM interval. The locus also has a pleiotropic effect on grain width (1.5 %) during late grain development that determines the relative magnitude of the grain weight increase. Positive NILs have increased maternal pericarp cell length, an effect which is independent of absolute grain length.
- These results provide direct genetic evidence that pericarp cell length affects final grain size and weight in polyploid wheat. We propose that combining genes which control distinct biological mechanisms, such as cell expansion and proliferation, will enhance crop yields.

## Introduction

By 2050, it is predicted that the human population will have exceeded nine billion people (United Nations, 2015). This is driving an increased demand for food production which is exacerbated by the use of crops for fuel and animal feed, and the pressures on agricultural systems resulting from climate change. With land for agricultural expansion being limited, increasing crop yields provides a sustainable route towards meeting this demand. However, rates of yield increase have slowed in recent years and are currently insufficient to achieve the estimated doubling in crop production that will be required by 2050 (Tilman *et al,* 2011; Ray *et al,* 2013). Projections show that increasing productivity on existing farmlands would increase the available food supply and lower prices, significantly reducing the number of people at risk of hunger globally (Rosegrant *et al,* 2013). With one in nine people currently living under food insecurity (FAO *et al,* 2015), it is urgent that we identify ways to increase crop yields.

Final crop yield is a complex quantitative trait strongly influenced by interacting genetic and environmental factors. For cereal crops, seed/grain weight (measured as thousand grain weight (TGW)) is a major yield component and is more stably inherited than final yield itself (Kuchel *et al,* 2007). Grain weight is largely defined by the size of individual grains and the morphometric components of grain area, length and width. A number of genes controlling these traits have been cloned from major grain weight quantitative trait loci (QTL) in rice (Fan *et al,* 2006;Song *et al,* 2007;Weng *et al,* 2008;Wang *et al,* 2012). For example, GW2, a RING-type E3 ubiquitin ligase acts as a negative regulator of cell division and was identified as the gene underlying a major QTL for rice grain width and weight (Song *et al,* 2007). These studies, in addition to those in model species, have shown that seed size is controlled through diverse mechanisms and genetic pathways (reviewed in Xing & Zhang, 2010; Li & Li, 2015). In Arabidopsis, the *AINTEGUMENTA* (*ANT*) transcription factor increases seed size through increased cell proliferation (Mizukami & Fischer, 2000), whilst the *APETELA2* (*AP2*) transcription factor regulates seed size by limiting cell expansion (Ohto *et al,* 2005). Other genes include those involved in phytohormone biosynthesis and signalling (Riefler *et al.* 2006; Schruff *et al.,* 2006; Jiang *et al.,* 2013) and G-protein signalling pathways (Huang *et al.,* 2009). Interestingly, many of these genes have been shown to act maternally (reviewed in Li &Li, 2015) and it has been proposed that the seed coat/pericarp (a maternal tissue) sets an upper limit to the final size of the seed/grain (Adamski *et al.,* 2009; Hasan *et al.,* 2011; Xia *et al.,* 2013).

Despite these advances, our understanding of the control of grain size is more limited in important crop species such as wheat (*Triticum aestivum*). Wheat provides around 20% of the calories consumed by humans and more protein globally than all types of meat combined (FAO, 2017). Many QTL for grain weight and, more recently, individual grain size/shape components have been identified in wheat (Breseghello & Sorrells, 2007; Gegas *et al.,* 2010; Simmonds *et al.,* 2014; Farre *et al.,* 2016; Kumar *et al.,* 2016). However, no mechanistic insight has been provided for these QTL, few have been validated (Simmonds *et al.,* 2014) and as yet, none have been cloned.

A major challenge to validate and define the mechanisms governing grain weight QTL in polyploid wheat has been that their effects are often subtle compared with QTL identified in diploid species such as rice (Uauy, 2017). One explanation is that wheat has a more limited capacity for increasing grain size than rice. An alternative, and more likely, scenario is that the effect of variation in an individual gene is masked by functional redundancy from homoeologous gene copies (Borrill *et al.,* 2015); bread wheat is a hexaploid species with three homoeologous genomes (A, B and D) that share between 96–98% sequence similarity across genes (Krasileva *et al.,* 2013). In addition, the size (17 Gb) and highly repetitive nature of the wheat genome has meant that, until recently, the genomic resources available in wheat have been limited. However, in the last few years there has been a radical change in the wheat genomics landscape with resources now including complete genome sequences and high quality gene models (IWGSC RefSeq v1.0; IWGSC, 2014; Clavijo *et al.,* 2017), transcriptomic databases (Pearce *et al.,* 2015b; Borrill *et al.,* 2016), high density single nucleotide polymorphism (SNP) arrays (Wang *et al.,* 2014; Winfield *et al.,* 2016) and exome-sequenced mutant populations (Krasileva *et al.,* 2017).

In this study, we identified a stable and robust QTL for grain weight in hexaploid wheat, which is driven by an increase in grain length. The QTL affects cell expansion in the grain and acts to increase the length of cells in the pericarp (maternal seed coat). We genetically mapped the effect to an interval on chromosome 5A, and used the latest wheat genome sequences and gene models to define the genes within the physical space. This detailed characterisation of the QTL provides direct genetic evidence that pericarp cell expansion affects final grain size, offering new insights into the mechanisms controlling grain weight in polyploid wheat.

## Methods

### Plant material

A doubled haploid (DH) mapping population was developed from the cross between two UK hexaploid winter wheat cultivars, ‘Charger’ and ‘Badger’. The population was created using the wheat x maize technique from F_1_ plants (Laurie & Bennett, 1988) and comprised 129 individuals, 92 of which were genotyped and used for evaluation. The 5A QTL was validated with the development of near isogenic lines (NILs). Two DH lines (CB53 and CB89) homozygous for the positive Badger loci across the complete linkage group were crossed to Charger and heterozygous F_1_ plants were backcrossed to the Charger recurrent parent for four generations (BC_4_). Heterozygous plants were selected at each generation using markers *Xgwm293* and *Xgwm186.* After BC_2_ and BC_4_, heterozygotes were self-pollinated and NILs homozygous for the alternative alleles across the interval were extracted (BC_2_F_2_ and BC_4_F_2_). In total 10 BC_2_ NILs were generated, six of which carried the *Xgwm293* to *Xgwm186* Badger positive interval. An additional twelve BC_4_ NILS were generated from the two DH lines (six Badger and six Charger interval).

Two representative BC_4_ NILs with alternative haplotypes were genotyped with the 90K iSelect array (Wang *et al.,* 2014) to confirm the introgression and identify additional segregating genomic regions. Recombinant BC_4_F_2_ plants between the flanking markers were also selected and self-pollinated for the development of homozygous BC_4_F_3_ recombinant inbred lines (RILs). Screening 170 plants with flanking markers *Xgwm293* and *Xgwm186* yielded 60 recombinants within the interval, defining a genetic interval of 17.65 cM.

### Genetic map construction and QTL analysis

The Charger x Badger genetic map was developed using simple sequence repeat (SSR) markers. From 650 SSR’s tested, 239 from JIC*/psp* (Bryan *et al.,* 1997;Stephenson *et al.,* 1998), IPK Gatersleben/*gwm*/*gdm* (Roder *et al.,* 1998; Pestsova *et al.,* 2000), Wheat Microsatellite Consortium/*wmc* (http://wheat.pw.usda.gov/ggpages/SSR/WMC/), Beltsville Agricultural Research Station*/barc* (Song *et al.,* 2005) and INRA/*cfa/cfd* (Guyomarc’h *et al.,* 2002) collections were polymorphic between parental lines. Consensus maps (Somers *et al.,* 2004) were used to select 212 SSR markers which maximised genome coverage with an approximate marker density of one SSR every 20 cM. In addition, nine sequence-tagged microsatellite profiling (STMP) markers (Hayden & Sharp, 2001) were incorporated into the map. To increase marker density, 75 Kompetitive Allele Specific Primers (KASP) markers were utilised. Markers with assigned chromosome locations (Allen *et al,* 2011) were targeted to fill gaps in the genetic map.

DNA extractions and genotyping procedures were performed as in Simmonds *et al.,* (2014). Likewise, map construction, QTL detection and multi-trait multi-environment (MTME) analysis was conducted as in Simmonds *et al.* (2014). Significant QTL effects were detected above a 2.5 log-of-odds (LOD) threshold.

SSR and KASP markers used in the QTL analyses were positioned with respect to the newly released Chinese Spring sequence through a BLAST search of 100 to 300 bp encompassing each SNP against the IWGSC RefSeq v1.0 (https://wheat-urgi.versailles.inra.fr/Seq-Repositorv/Assemblies). In most cases, the order on the reference sequence agreed with the genetic order in the Charger x Badger population. For discrepancies, we used the genetic position to order markers. In cases of no hits to the RefSeq v1.0 assembly, we inferred a physical position based on the two closest markers and the relative distance of all three markers based on their centiMorgan positions. Similarly, physical positions of all iSelect SNPs were obtained using BLAST to align the surrounding sequence (201 bp) to the RefSeq v1.0 assembly. TGACv1 gene models were positioned on RefSeqv1.0 with GMAP (Wu & Watanabe, 2005) using best hit position and 95% minimum similarity cut-off (David Swarbreck and Gemy Kaithakottil; Earlham Institute).

### Field evaluation and phenotyping

The DH population was evaluated in the field in a randomised complete block design with three replications at five sites (Norwich and Sandringham, England; Balmonth, Scotland; Bohnshausen, Germany; and Froissy, France (Simmonds *et al.,* 2014)). The experiments were grown for three years (2001–2003) at Norwich and Sandringham, and two years (2002–2003) at the other three sites. The field trials were sown in large-scale yield plots (1.1□×□6 m) and treated with standard farm pesticide and fertiliser applications to reproduce commercial practise. All trials were sown by grain number for comparable plant densities per plot (275 seeds*m^−2^). Plots were measured for final plot yield after adjustment for plot size, and TGW was calculated by counting and weighing 100 seeds from each plot.

The NILs were evaluated at Norwich in 2012 and 2013 (10 BC_2_ NILs), 2014 (12 BC_4_ NILs) and 2015 and 2016 (4 BC_4_ NILs), while BC_4_ RILs were analysed in 2014–2016. For both NIL and RIL experiments, a randomized complete block design was used with five replications. NILs were grown in large-scale yield plots (1.1+×+6 m), whereas RILs were grown in 1.1 × 1 m plots in 2014 and large-scale yield plots in 2015 and 2016. Final grain yield (adjusted by plot size and moisture content) was determined for NILs across the five years. Developmental traits were also measured for NILs in 2012–2016, although not all traits were measured in each year (Supporting Information Table S1). For all NILs (2012–2016) and RILs (2014–2016), grain morphometric measurements (grain width, length, area) and TGW were recorded on the MARVIN grain analyser (GTA Sensorik GmbH, Germany) using approximately 400 grains obtained from the harvested grain samples. For all NILs (2012–2016), ten representative spikes per field plot were also measured for spike yield components (spikelet number, number of viable spikelets, spike length, grain number per spike, spike yield and seeds per spikelet), TGW and grain morphometric parameters. The data from the ten representative spikes was consistent with the whole plot values.

### Grain developmental time courses

The BC_4_ NILS grown in 2014–2016 were used for the grain developmental time courses. Two Charger (5A-) and two Badger (5A+) NILs were used, and the same NILs were used in all three years. We tagged 65 ears per NIL across each of four blocks in the field at full ear emergence (peduncle just visible) to ensure sampling at the same developmental stage. Ten spikes per NIL, per block, were sampled at each of five (2014) or six (2015–2016) time points. 2014 time points included 4, 8, 12, 18 and 27 days post anthesis (dpa). 2015 time points included anthesis (0 dpa), 4, 7, 12, 19 and 26 dpa. 2016 time points included 0, 3, 8,10,15, 21 dpa. Ten grains were sampled from each spike from the outer florets (positions F1 and F2) of spikelets located in the middle of the spike. Grains were weighed to obtain fresh weight, assessed for morphometric parameters (grain area, length and width) on the MARVIN grain analyser and then dried at 37 °C to constant weight (dry weight). For each block at each time point, a total of ~100 grains were sampled (10 spikes × 10 grains) per NIL However, for the statistical analysis the average of each NIL within each block was used as the phenotypic value as the individual grains and spikes were considered as subsamples.

### Cell size measurements

One representative 5A- and 5A+ BC_4_ NIL was used for cell size measurements. We selected mature grains from three blocks of the 2015 harvest samples based on a variety of criteria. For each NIL, we selected nine grains of average grain length from the whole harvest sample from each block (groups 5A-/5A+ average). For the 5A- NIL, an additional nine grains were selected that had grain lengths equivalent to the average of the 5A+ NIL sample (5A- large). For the 5A+ NIL an additional nine grains were selected that had grain lengths equivalent to the average of the 5A- NIL sample (5A+ small). We also selected grains of average length from three blocks of the 2016 harvest (nine grains were selected from each block per genotype). Grains were stuck crease-down on to 12.5mm diameter aluminium specimen stubs using 12 mm adhesive carbon tabs (both Agar Scientific), sputter coated with gold using an Agar high resolution sputter coater and imaged using a Zeiss Supra 55 SEM. The surface (pericarp) of each grain was imaged in the top and bottom half of the grain, with images taken in at least three positions in each half. All images were taken at a magnification of 500x. Cell length was measured using the Fiji distribution of ImageJ (Schindelin *et al.,* 2012) (Supporting Information Fig. S1). Cell number was estimated for each grain using average cell length/grain length. For the statistical analyses, we considered the average cell length of each individual grain as a subsample within the block.

### Statistical analysis

DH lines homozygous across the genetic interval for the two major QTL, *Qtgw-cb.2B* (*Xgwm259- Xstm119tgag*) and *Qtgw-cb.5A* (.*Xgwm443-XBS00000435*) were classified by genotype. Using this classification, general linear model analyses of variance (ANOVA) were performed for TGW incorporating environment and year as factors for each individual QTL, and for lines with both increasing alleles compared with those with neither. Pearson’s correlation coefficient was calculated to assess the correlation between yield and TGW. All analyses performed on DH lines was carried out using Minitab v17.3.1 (Minitab Inc.).

The NILs and RILs were evaluated using two-way ANOVAs, with the model including the interaction between environment and the 5A QTL. RIL groups were assigned as having a Charger or Badger like grain length phenotype using a post hoc Dunnett’s test to compare with C- and B-control groups. Similarly, two-way ANOVAs, including genotype and block, were conducted for the developmental time courses and cell size measurements. Analyses were performed using GenStat 15^th^ edition (VSN International) and R v3.2.5.

## Results

### A QTL on chromosome 5A is associated with increased grain weight

A genetic map was developed for the Charger × Badger DH population comprising of 296 polymorphic molecular markers. Linkage analysis resulted in 32 linkage groups which were assigned to 21 chromosomes, covering a genetic distance of 1,296 cM. The only chromosome with no marker coverage was 6D.

QTL analysis identified two regions with consistent variation for TGW, chromosomes 2B (*Qtgw-cb.2B*) and 5A (*Qtgw-cb.5A*), based on the mean LOD score across environments (Fig. 1) and colocalization of significant QTL (Supporting Information Table S2). *Qtgw-cb.2B* was identified in 7 of the 12 site/year environments, providing a mean of 11% of the explained variation when significantly expressed and a mean additive effect of 1.26 g/1000 grains, with Charger providing the increasing allele. The peak LOD for the QTL was located at markers *Xgwm148* and *Xgwm120* depending on the environment. *Qtgw-cb.5A* was also significant at 7 of the 12 environments and accounted for 15.5% of the phenotypic variation with a mean additive effect of 1.6 g/1000 grains. The peak for *Qtgw-cb.5A* was defined by markers *Xgwm293* (20.6 cM) and *Xbarc180* (25.3 cM; Fig. 2a), with Badger providing the increasing allele.

**Figure 1:**
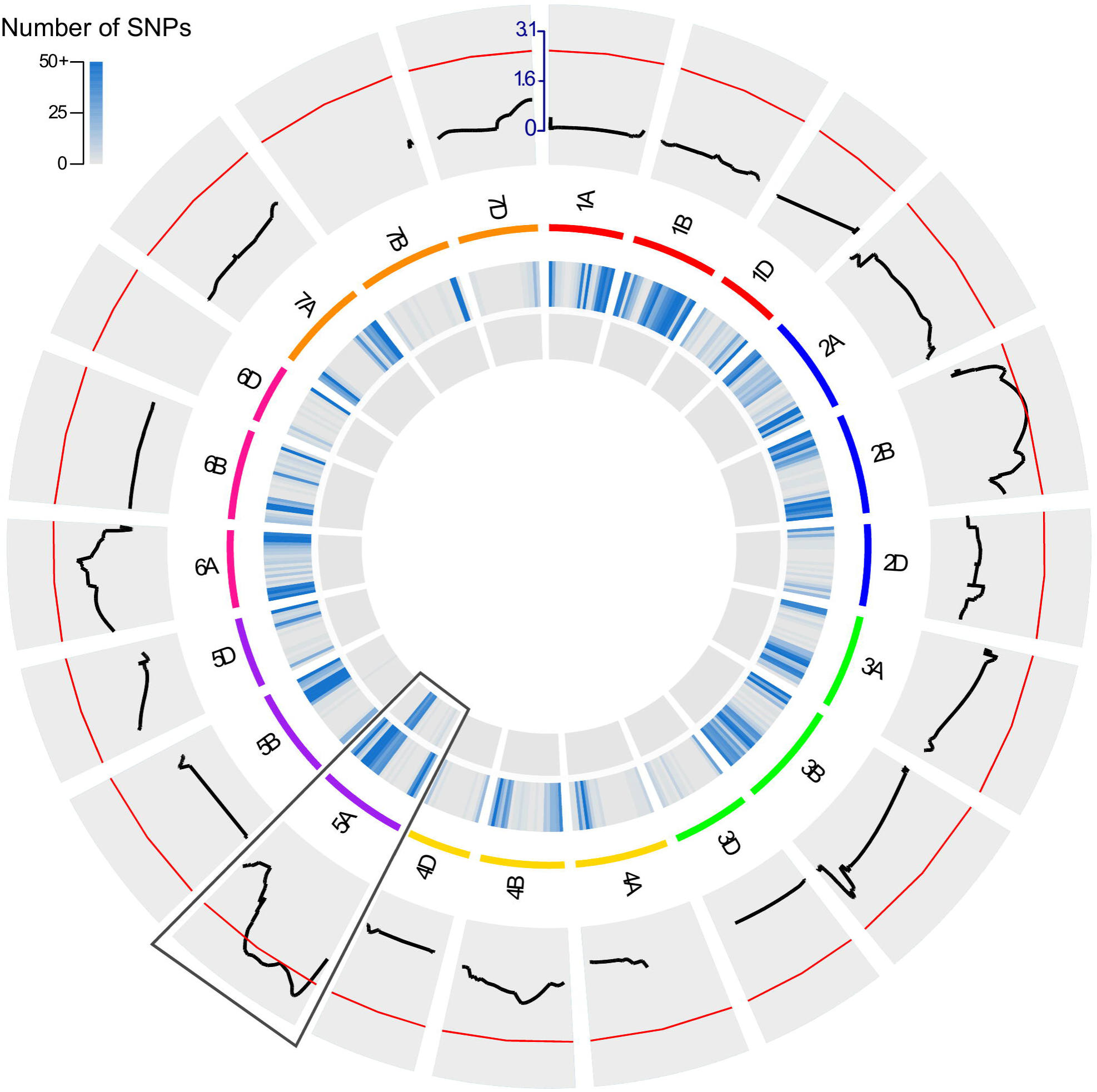
QTL analysis and NIL development. Circos diagram showing the whole genome quantitative trait loci (QTL) scan and single nucleotide polymorphism (SNP) variation. Outer track is the mean log-of-odds (LOD) score for thousand grain weight (TGW) across all environments measured. The red line shows a LOD threshold of 2.5. Wheat chromosome groups are represented in different colours beneath the QTL scans. The most significant and stable QTL identified was on chromosome 5A (boxed segment). Inner tracks correspond to heatmaps representing the number of iSelect SNPs in 30 Mb windows showing variation between Charger and Badger, parents of the doubled haploid population (outer); ora representative pair of 5A-/5A+ NILs (innermost). Physical positions of all markers (including those used in the QTL scan and iSelect markers) were determined using the IWGSC RefSeq v1.0 sequence.

**Figure 2:**
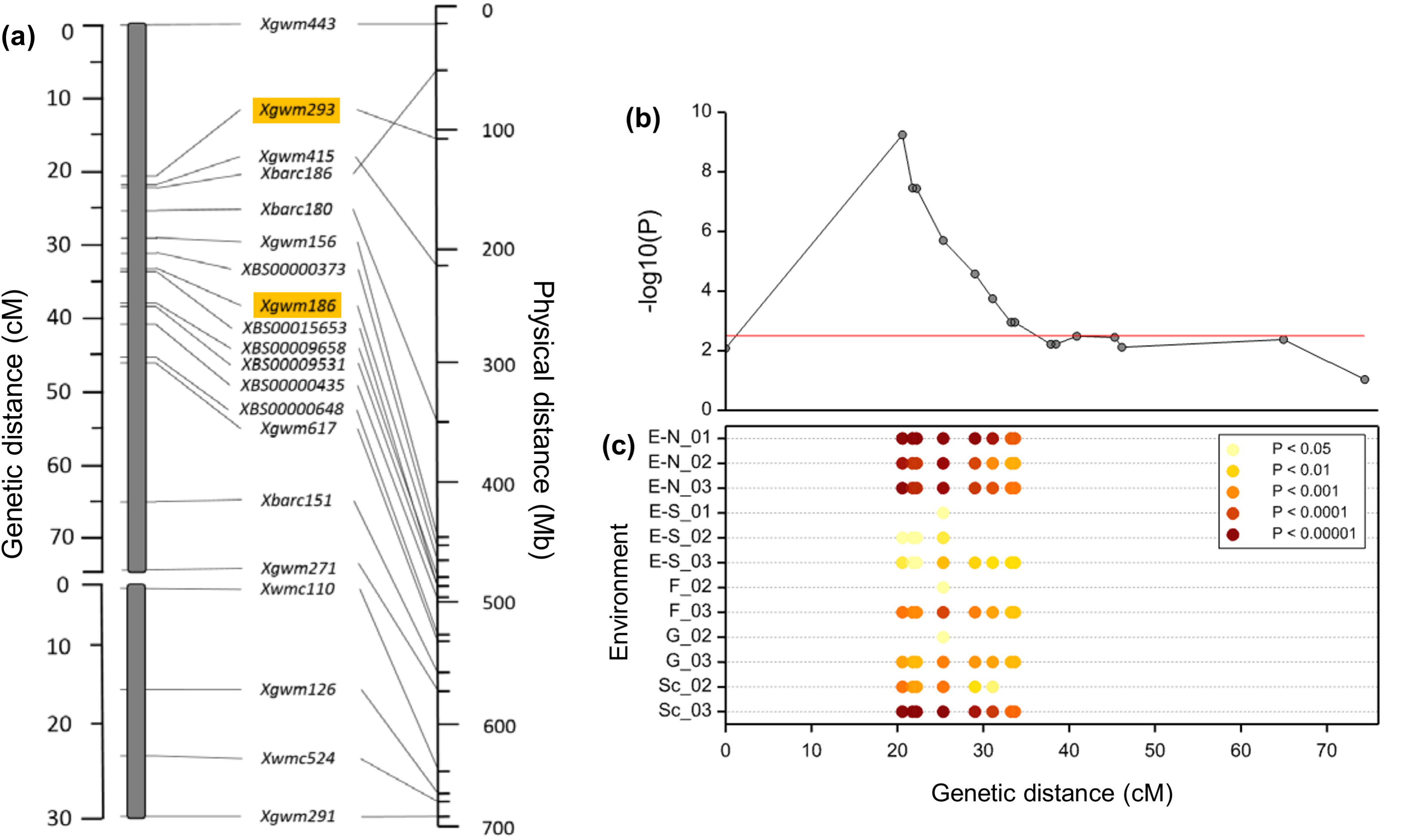
Chromosome 5A genetic/physical map and TGW MTME analysis. (a) Genetic and physical map of chromosome 5A. The left-hand side represents the genetic map, comprised of two linkage groups with calculated distances between markers in cM (Linkage group 1: 0–74.4 cM, Linkage group 2: 0–30.2 cM). The right-hand side represents the physical map according to the Chinese Spring IWGSC RefSeq v1.0 sequence. Markers highlighted in orange indicate those used for NIL development, (b) Multi-trait multi-environment (MTME) QTL analysis of the 5A QTL for thousand grain weight (TGW) across Linkage group 1. The red line indicates a log-of-odds (LOD) threshold of 2.5. (c) Markers with significant additive effects are shown for each environment for those markers above the LOD threshold in (b). The intensity of the colour (yellow to brown) indicates the level of the significance as indicated by the legend. ΕΝ = England-Norwich, E-S = England, Sandringham, F = France, G = Germany, Sc = Scotland.

Analysis of DH lines homozygous across the wider QTL regions for both *Qtgw-cb.2B* (*Xgwm259- Xstm119tgag)* and *Qtgw-cb.5A* (*Xgwm443-XBS00000435*) demonstrated that the increasing alleles of each individual QTL provided a significant 4.1% and 5.5% increase in TGW (P < 0.001), respectively. DH lines containing both QTL (n=9) produced a 10% increase (P < 0.001) over lines with neither (n=10), suggesting *Qtgw-cb.2B* and *Qtgw-cb.5A* are additive when combined.

There was a significant correlation (P < 0.001) between grain yield and TGW across all datasets, however significant QTL were only co-located for both traits in France 2003 (2B) and England-Norwich 2002/Scotland 2002(5A) (Supporting Information Table S3). This suggests that although TGW was an important component regulating yield in this DH population, it was also influenced by other yield components. As *Qtgw-cb.5A* had a larger mean additive effect and accounted for more of the phenotypic variation than *Qtgw-cb.2B,* we selected *Qtgw-cb.5A* for further analyses.

### Multi-Trait Multi-Environment (MTME) analysis defines *Xgwm293* as the peak marker of *Qtgw-cb.5A*

MTME analysis was conducted on chromosome 5A for both TGW and grain yield. For TGW, markers above the significance threshold (LOD >2.5) ranged from *Xgwm293* (20.6 cM) to *XBS00015653* (33.7 cM) with the peak being at *Xgwm293* (Fig. 2b,c). At least one of the markers within the identified region was significant at each of the twelve environments, with Badger always providing the beneficial alleles. For yield, MTME analysis identified a significant QTL in the *Qtgw-cb.5A* region, with the peak marker (*Xgwm293*) being the same as for TGW. Significant increases in the additive effect of Badger were observed in seven environments (Supporting Information Fig. S2), contrasting to only two in the previous single-environment analysis. It is worth noting that in two environments (England-Sandringham 2001 and 2003), the alternative parent Charger had a borderline significant effect on yield in the MTME analysis. Taken together, these results suggest that the Badger *Qtgw-cb.5A* interval is associated with a consistent effect on TGW across environments which often, but not always, translates into a yield benefit.

### Near isogenic lines (NILs) differing for *Qtgw-cb.5A* show a 6.9% difference in TGW

To independently validate and further investigate the effect of *Qtgw-cb.5A* (referred to hereafter as 5A QTL) on TGW, BC_2_ and BC_4_ NILs differing for the QTL region were developed using markers *Xgwm293* and *Xgwm186* and Charger as the recurrent parent. Pairs of BC_4_ NILs carrying the Charger (5A-) or Badger (5A+) segment were genotyped using the iSelect 90K SNP array (Wang et al, 2014) and found to be 97.2% similar, only showing variation in 221 markers across the 5A QTL, compared with 7,973 SNPs between the parents (Fig. 1, inner tracks). These NILs therefore provide a valuable resource for specifically studying the effects of the 5A QTL in more depth.

Across five years of replicated field trials 5A+ NILs showed an average increase in TGW of 6.92% (P < 0.001) ranging from 4.00 to 9.28% (Table 1), and significant in all years. The difference in TGW was associated with a yield increase of 1.28% in 5A+ NILs across all years, although this effect was not significant (P = 0.093). The effect varied across years with a significant yield increase of 2.17% (P = 0.046) in 2014 and non-significant effects of between 0.02 to 1.72% in the other four years. The positive effect of the QTL on yield was similarly subtle in the DH population as described previously.

**Table 1:** Mean Thousand Grain Weight (TGW), yield and grain morphometric parameters of 5A NILs

| Year | Genotype | TGW (g) | Yield (kg/plot) | Grain area (mm<sup>2</sup>) | Grain length (mm) | Grain width (mm) |
| --- | --- | --- | --- | --- | --- | --- |
| 2012 | 5A- | 38.027 | 4.408 | 18.755 | 6.625 | 3.475 |
|  | 5A+ | 41.554 | 4.437 | 19.930 | 6.900 | 3.557 |
|  |  | 9.28%*** | 0.66%<sup>NS</sup> | 6.26%*** | 4.15%*** | 2.35%** |
| 2013 | 5A- | 40.772 | 6.157 | 19.969 | 6.705 | 3.674 |
|  | 5A+ | 43.544 | 6.159 | 20.979 | 6.963 | 3.727 |
|  |  | 6.80%*** | 0.02%<sup>NS</sup> | 5.06%*** | 3.86%*** | 1.44%*** |
| 2014 | 5A- | 47.368 | 6.495 | 21.493 | 6.798 | 3.930 |
|  | 5A+ | 50.729 | 6.636 | 22.579 | 7.063 | 3.979 |
|  |  | 7.09%*** | 2.17%* | 5.05%*** | 3.90%*** | 1.25%** |
| 2015 | 5A- | 42.734 | 7.582 | 18.044 | 6.426 | 3.479 |
|  | 5A+ | 46.201 | 7.712 | 19.293 | 6.730 | 3.554 |
|  |  | 8.11%*** | 1.72%<sup>NS</sup> | 6.93%*** | 4.72%*** | 2.16%*** |
| 2016 | 5A- | 49.292 | 5.974 | 19.829 | 6.580 | 3.735 |
|  | 5A+ | 51.266 | 6.064 | 20.610 | 6.816 | 3.745 |
|  |  | 4.00%* | 1.50%<sup>NS</sup> | 3.94%** | 3.58%*** | 0.27%<sup>NS</sup> |
| Overall | 5A- | 43.639 | 6.123 | 19.618 | 6.627 | 3.659 |
|  | 5A+ | 46.659 | 6.201 | 20.678 | 6.894 | 3.712 |
|  |  | 6.92%*** | 1.28%<sup>NS</sup> | 5.41%*** | 4.04%*** | 1.45%***<sup>1</sup> |

<sup>1</sup> %s indicate amount gained in 5A+ NILs compared with 5A- NILs. Superscripts indicate significance determined by ANOVA for either each year, or across all years (final row). ie. NS = Non-significant, \* = P < 0.05, \*\* = P < 0.01, \*\*\* = P < 0.001. 2012-13 = BC<sub>2</sub>-NILs, 2014-16 = BC<sub>4</sub>-NILs.

We measured the NILs for a series of spike yield component traits to determine possible pleiotropic effects associated with the 5A+ TGW effect. Within most years, there was no significant effect of the 5A+ allele on spike yield components such as spikelet number, seeds per spikelet or grain number per spike (Supporting Information Table S4). However, when all years were analysed together, we observed a significant reduction in grain number (−3.55%, P = 0.04) and seeds per spikelet (−3.37%, P = 0.015) associated with the 5A+ interval. This statistical significance was driven by a particularly strong negative effect in 2016 as grain number and seeds per spikelet were non-significant in the preceding four seasons (2012-15). Overall, however, the 5A+ interval is associated with a consistent small decrease in these spike yield components.

Taking into account the 6.92% effect of the 5A+ QTL on TGW and the tendency for decreases in some spike yield components, the overall spike yield increased by 2.33% (P = 0.032) across the five years. However, similar to grain number and seeds per spikelet, the statistical significance is driven by a single year (2014) despite overall positive effects in another three years (2012, 2013, and 2015). We also measured tiller numbers and found a significant reduction of 4 tillers per m in the 5A + NILs across two years (P = 0.008) (Supporting Information Table S1). No effect was seen for spikelet number and additional phenology traits (Supporting Information Table S4). Taken together, these results suggest that the 5A+ interval has a consistent positive effect on TGW and that the effects on yield are modulated by a series of smaller compensating negative effects on yield components such as grain number, seeds per spike and tiller number.

### The TGW increase in 5A+ NILs is primarily due to increased grain length

TGW is determined by individual components including physical parameters such as grain length and width. To understand the relative contribution of these components to the increase in TGW, NILs were assessed for these grain morphometric parameters (length, width and area) using a 2D imaging system (Table 1). 5A+ NILs had significantly increased grain length (P < 0.001), width (P < 0.001) and area (P < 0.001) compared to 5A- NILs across all years with the exception of width in 2016. On average, the 5A+ QTL increased grain length by 4.04% (P < 0.001), ranging from 3.58 to 4.72% (P < 0.001 in all years). The effect on width was smaller, averaging 1.45% (P < 0.001; range 0.27 to 2.35%) and significant in four out of five years (Table 1). The effects on length and width combined to increase grain area by an average of 5.41% (P < 0.001), significant in all five years. These results were based on combine harvested grain samples and were also confirmed in ten representative single ear samples taken before harvest. TGW of the ten spikes correlated strongly with the whole plot samples (r =0.84, P < 0.001) and showed a similar difference between NILs (6.00%, P < 0.001; Supporting Information Table S4). Across datasets, the effect of the 5A+ QTL on grain length was more than twice the size of the effect on grain width. This fact, together with the more consistent effect on grain length across years (Coefficient of variation length = 10.6%; width = 55.3%; TGW = 27.8%) suggests that the increase in grain length is the main factor driving the increase in grain area and TGW.

We compared the distribution of grain length and width using data from individual seeds to determine whether the QTL affects all grains uniformly. Violin plots for length showed variation in distribution shape among years (Fig. 3). However, within years the 5A- and 5A+ grain length distributions were very similar in shape, suggesting that the QTL affects all grains uniformly and in a stable manner across the ear and within spikelets. In all years, the 5A+ grain length distributions were shifted higher than the 5A- NILs with an increase in longer grains and fewer shorter grains, in addition to the higher average grain length (Fig. 3). Grain width distributions were also very similar in shape within years, but had a less pronounced shift between NILs (Supporting Information Fig. S3) consistent with the overall smaller effect of the 5A QTL on grain width.

**Figure 3:**
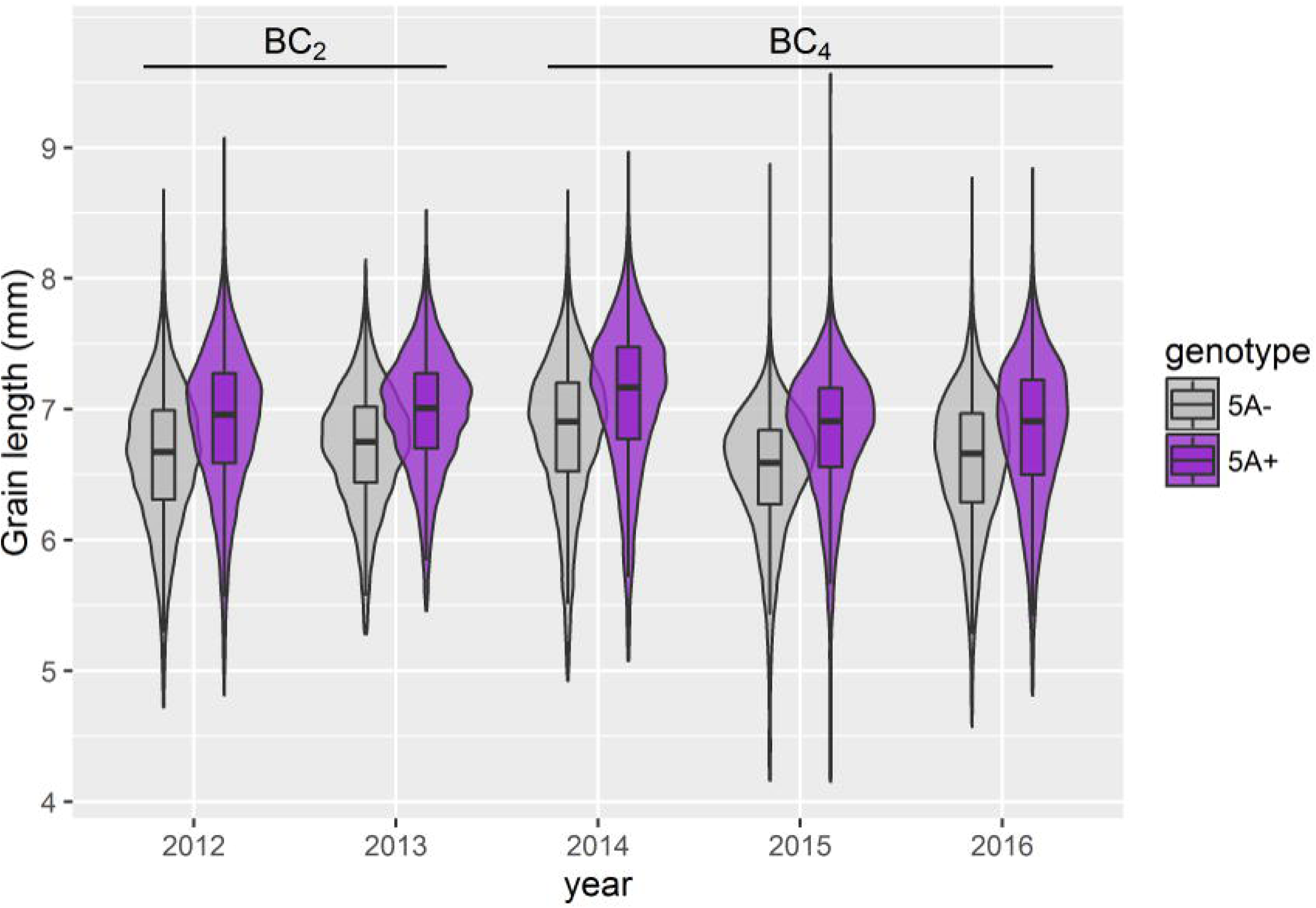
Distribution of grain length of NILs from whole plot samples. Violin plots showing the distribution of individual seed measurements of grain length across the five field experiments of BC_2_ (2012–2013) and BC_4_ (2014–2016) near isogenic lines (NILs). Purple = 5A+ NILs, grey plots = 5A-NILs. All within year comparisons between NILs were significant (P < 0.001).

### The 5A QTL region acts during grain development to increase grain length

To determine when differences in grain morphometric parameters between NILs are first established, we conducted grain development time courses of two 5A- and two 5A+ BC_4_ NILs. Grains were sampled in 2014, 2015 and 2016 from field plots at anthesis and at five further time points across grain development until the difference in grain size had been fully established.

Data from 2015 is shown in Fig. 4 as a representative year (samples taken at anthesis (0 dpa), 4, 7,12,19 and 26 dpa). The first significant difference in grain length was observed at 12 dpa with 5A+ NILs having 1.5% longer grains than 5A- NILs (p =0.034). This effect increased to 4.4 % at 19 dpa (P < 0.001) and was maintained at 26 dpa (4.5 % increase, P < 0.001; Fig. 4a). No significant effects on grain width were observed until 26 dpa when 5A+ NILs increased grain width by 1.7 % (P = 0.015; Fig. 4b). Significant differences in grain area were detected at 19 dpa (5.7 % increase; P < 0.001; data not shown) and this difference was maintained at the final time point 26 dpa (6.1 %, P < 0.001). By the final time point 5A+ NILs also had significantly heavier grains (3.7%, P = 0.01; Fig. 4c). These effects were all consistent with the grain size and weight differences observed in mature grains in 2015 (Table 1) and were also observed in 2014 and 2016 (Supporting Information Figs. S4,S5). The fact that the effects on width, area and weight are all after the first significant difference on grain length in all three years further supports grain length as the main factor driving the increase in grain weight.

**Figure 4:**
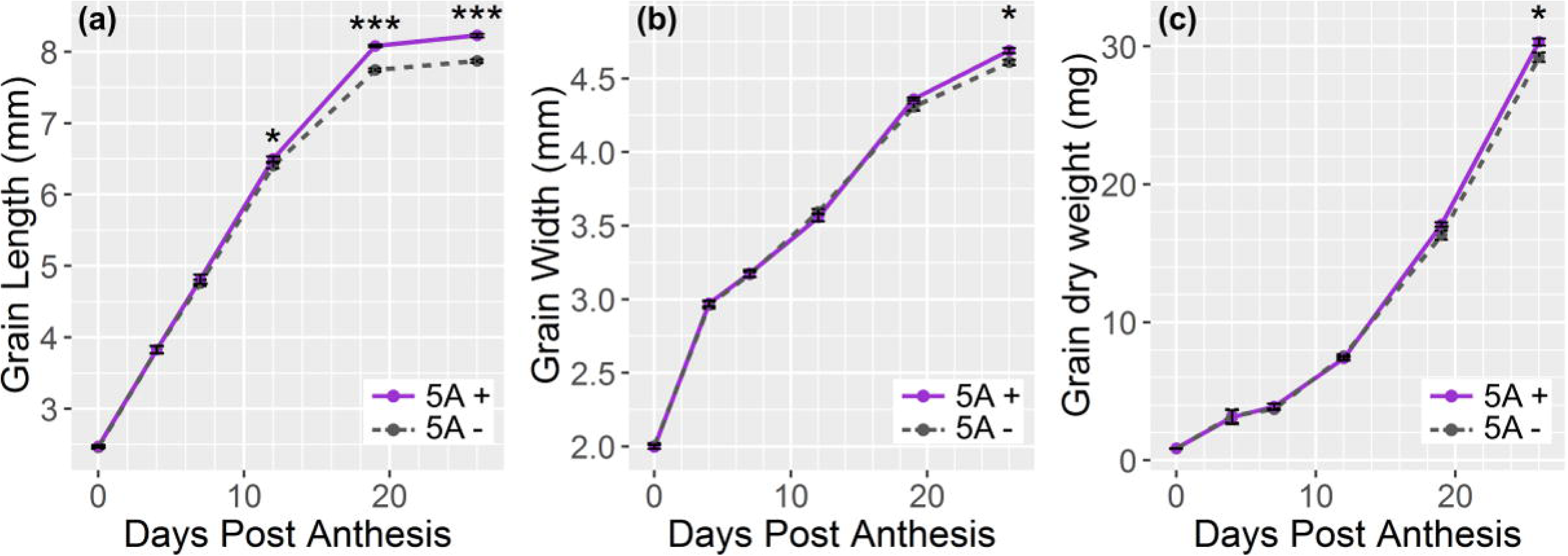
Grain development time course of 5A- and 5A+ NILs. Grain length (a), grain width (b) and grain dry weight (c) of 5A- (grey, dashed line) and 5A+ (purple, solid line) BC_4_ near isogenic lines (NILs) during grain development with samples taken at anthesis (0 days post anthesis (dpa)), 4, 7,12,19 and 26 dpa in 2015 field trials. ^*^ = P < 0.05, ^**^ = P < 0.01, ^***^ = P < 0.001. Error bars show standard error of the mean.

### 5A+ NILs have increased pericarp cell length independent of absolute grain length

We used scanning electron microscopy (SEM) to image pericarp cells and determine cell size of BC_4_ 5A- and 5A+ grains. Mature grains from the 2015 field experiment were selected from a 5A- and 5A+ NIL pair based on their grain length and using a variety of criteria to allow for distinct comparisons (Fig. 5). First, we compared grains of average length from the 5A- and 5A+ NIL distributions (Fig. 5a). We found that average 5A+ grains had an 8.33 % significant increase in mean cell length (P = 0.049) compared to average 5A- grains and that this was reflected in a shift in the whole distribution of 5A+ cell lengths (Fig. 5a). Next, we compared cell lengths in grains of the same size from 5A- and 5A+ NILs. We selected relatively long grains from the 5A- NIL distribution (Fig. 5b; orange) that had the same grain length as the average 5A+ grains. This comparison showed that 5A+ grains still had longer cells (9.53%, P = 0.015) regardless of the fact that the grain length of the two groups were the same (6.8 mm; Fig. 5b). We also made the opposite comparison by selecting relatively short grains from the 5A+ NIL distribution (Fig. 5c; green) and comparing them with average 5A- grains. Similar to before, the 5A+ grains had longer cells (8.61%), although this effect was borderline non-significant (P = 0.053; Fig. 5c). Finally, a comparison of long 5A- grains and short 5A+ grains again showed that cells were longer in 5A+ grains (9.81%, P = 0.011), even though the 5A+ grains used in this comparison were 7.65% shorter than the 5A- grains. Within genotype comparisons of cell length between grains of different lengths showed no significant differences in mean cell length (Supporting Information Fig. S6). The results were confirmed in 2016 where average 5A+ grains had a 24.6 % significant increase in mean cell length compared to average 5A- grains (P < 0.001; Supporting Information Fig. S7). These results indicate that the 5A+ region from Badger increases the length of pericarp cells independent of absolute grain length. Using grain length and mean cell length to calculate cell number, we determined that the average length grains of both 5A- and 5A+ had the same number of cells in 2015. However, in 2016, 5A- NILs had significantly more cells than 5A+ NILs (Fig. S8).

**Figure 5:**
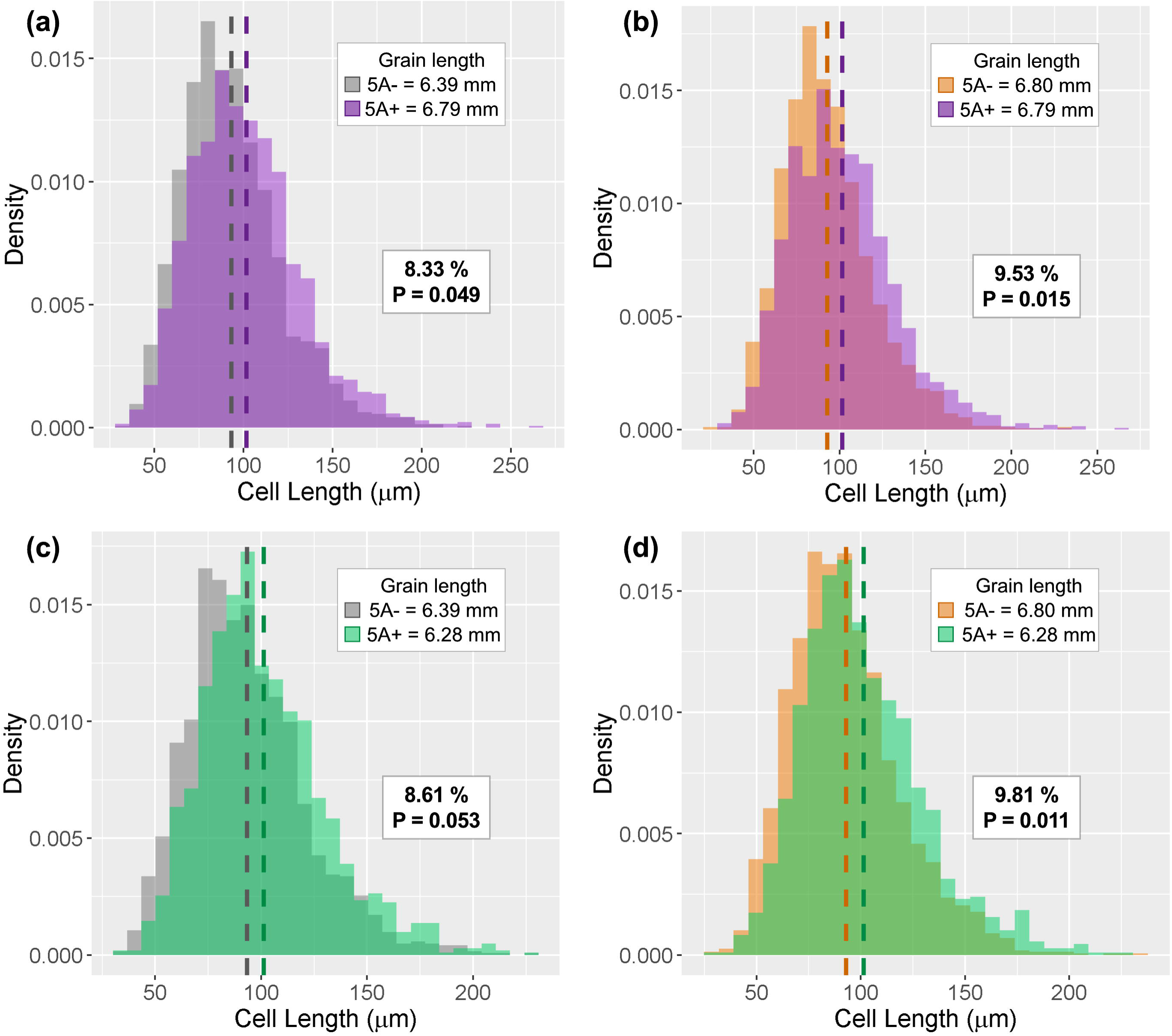
Comparisons of cell length between 5A- and 5A+ NILs. Density plots of cell length measurements from 27 grains per genotype group; dashed line represents the mean. “Grain length” insets show the average grain length of each group of grains used for measurements. The increase in cell length of 5A+ near isogenic lines (NILs) relative to cell length of 5A -grains is shown as a percentage along with the P values calculated using ANOVA to compare means of the two groups displayed, (a) Grains of average length from 5A- and 5A+ NILs, (b) average 5A+ grains and equivalent 5A- grains, (c) average 5A- grains and equivalent 5A+ grains, (d) long 5A- grains (length equivalent to average 5A+ grains) and short 5A+ grains (grain length equivalent to average 5A- grains).

### The grain length QTL maps to a 75 Mb / 4.3 cM genetic interval

We used a set of 60 homozygous RILs to map the grain length phenotype to a narrower genetic interval within the 5A QTL region (17.65 cM, 367 Mbp). KASP markers were developed for 25 additional SNPs between the two original QTL flanking markers (*Xgwm293* and *Xgwm186;* Fig. 6a) based on data from the iSelect genotyping of BC_4_ NILs and 820K Axiom Array genotyping of Charger and Badger (Winfield *et al.,* 2016). Based on the genotype of these 25 markers, 49 of the RILs were assigned to eleven distinct recombination groups represented as graphical genotypes in Fig. 6a. Control RILs were selected based on having either the Charger (5A-) or Badger (5A+) genotypes across the interval (C-control and B-control, respectively).

**Figure 6:**
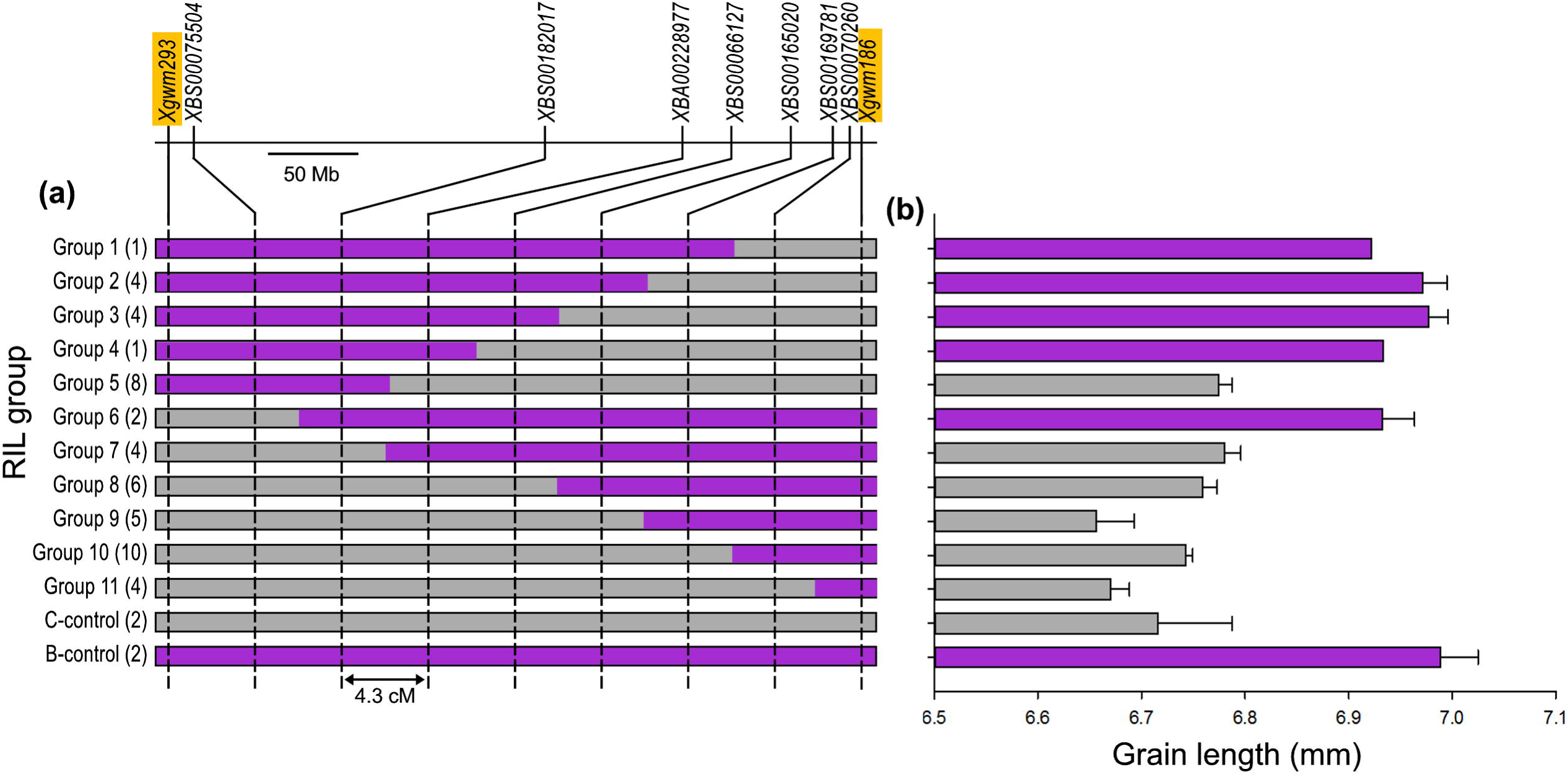
Grain length maps to a 4.3 cM interval on chromosome 5A. (a) Graphical genotypes of recombinant inbred lines (RIL) groups with the number of lines in each group shown in parentheses. RILs were grouped based on their genotypes defined by having either the Charger-like (grey) or Badger-like (purple) allele at each marker shown across the interval. Markers highlighted in orange indicate markers used for NIL development, (b) ANOVA adjusted mean grain length of RIL groups across all experiments. Bars are coloured based on a Charger- or Badger-like phenotype, determined by Dunnett’s test. Purple = Badger-like, grey = Charger - like. Error bars represent standard error of the mean.

RILs were phenotyped for grain length in three field seasons and we found significant differences between RIL groups (P < 0.001). The overall average grain length of the B-control group was 4.06 % higher than the C-control group (P < 0.001; Fig. 6b), consistent with the differences in grain length observed between the NILs (Table 1). Each RIL group was classified based on Dunnett’s tests to both control groups: for example, a RIL group was classified as Charger-like only if it was both significantly different to the B-control *and* non-significantly different to the C-control. Using this classification, we assigned unambiguously the eleven RIL groups to a parental type and genetically mapped the grain length phenotype between markers *XBS00182017* and *XBA00228977* (Fig. 6). This represents a genetic distance of 4.32 cM corresponding to a physical interval of 74.6 Mb in the Chinese Spring RefSeq v1.0 sequence.

This 74.6 Mb interval contains 811 TGACv1 gene models (Clavijo *et al.,* 2017) based on *in silico* mapping to the Chinese Spring reference (Supporting Information File S1). We analysed the expression profile of these genes on the wheat expVIP expression platform (Borrill et al., 2016) and found that 439 of these genes are expressed (>2 transcripts per million (tpm)) in at least one grain RNA-seq sample (n=147). The developmental time courses suggest that the 5A QTL acts at around 12 dpa and we found 405 of these transcripts expressed in grain samples taken at around this time (4–15 dpa, n = 59), with 298 genes expressed in the pericarp tissue (Pearce *et al.,* 2015a).

## Discussion

In this study we identified a stable and robust QTL associated with a 6.9 % increase in grain weight. This increase is driven by longer grains associated with increased pericarp cell length. In wheat and barley pericarp cell division decreases shortly after fertilization (2 to 6 days; Drea *et al.,* 2005; Radchuk *et al.,* 2011) and cell expansion plays the predominant role in increasing pericarp size during grain development. Our results are consistent with a role of the 5A gene on pericarp cell expansion given that significant differences in grain size are only observed twelve days after fertilization, once cell expansion has begun. However, we cannot discard an overlapping late effect on cell division given the conflicting results in final pericarp cell number between years.

Overall, our results suggest that the gene underlying this locus regulates, either directly or indirectly, cell expansion in the pericarp (seed coat), a mechanism that is known to be a key determinant of grain/seed size in several species. Some genes, such as expansins and XTH (xyloglucan endotransglucosylase/hydrolases), affect cell expansion directly by physically modifying or “loosening” the cell wall (reviewed in Cosgrove, 2005), and the expression of these enzymes has been associated with pericarp cell expansion in wheat and barley (Lizana *et al.,* 2010; Radchuk *et al.,* 2011; Munoz & Calderini, 2015). Other genes regulate pericarp/seed coat cell size indirectly, for example through the regulation of sugar metabolism and subsequent accumulation in the vacuole (Ohto *et al.,* 2005) and endoreduplication (Chevalier *et al.,* 2014). Our results provide direct genetic evidence that pericarp cell expansion affects final grain size and weight in polyploid wheat.

The maternal control of grain/seed size has been well documented in rice and Arabidopsis (Li &Li, 2015), as well as in wheat through physiological and genetic studies (Hasan *et al.,* 2011; Simmonds *et al.,* 2016). This can affect cell proliferation and/or cell expansion of maternal tissues, such as the wheat pericarp, both pre- and post-fertilisation (Garcia *et al.,* 2005; Adamski *et al.,* 2009; Ma *et al.,* 2016). For example, *GW2* in rice and its orthologue in Arabidopsis (DA?) affect grain/seed size through suppression of cell proliferation (Song *et al,* 2007; Xia *et al.,* 2013). Similarly in wheat, a knock-out mutant of the *GW2* orthologue has larger carpels than wild-type plants suggesting that the gene acts on maternal tissue pre-fertilisation (Simmonds *et al.,* 2016). The effect of the wheat *GW2* gene on cell size and number has not been determined however.

The direct assignment of the 5A effect to the maternal parent will require additional studies, including analysis of F_1_ hybrids from reciprocal crosses. These studies are not routinely performed in wheat given that the phenotypic variation between individual F_1_ grains often surpasses the relatively subtle phenotypic effects of most grain size QTL (usually less than 5% in wheat). The identification of a robust effect on pericarp cell length in this study, which is independent of the individual grain size, opens up a new approach to explore these parent-of-origin effects in polyploid wheat.

It has been proposed, in multiple species, that the size of the pericarp/seed coat determines final grain size by restricting endosperm growth (Calderini *et al.,* 1999; Adamski *et al.,* 2009; Hasan *et al.,* 2011). This is analogous to the way in which grain size in rice is limited by the size of the spikelet hull (Song *et al.,* 2005). Both the length (Lizana *et al,* 2010; Hasan *et al.,* 2011) and the width (Gegas *et al,* 2010; Simmonds *et al.,* 2016) of the pericarp have been proposed as key determinants of final grain weight in wheat. Our results provide genetic evidence for the importance of the maternal pericarp tissue and show that length is the underlying component for the 5A locus. Across three years, the difference in grain length between NILs was the first grain size component difference to be established. Only after this, did we observe any differences in grain width, weight or grain filling rate. These differences in grain length were extremely consistent across years (despite average TGW values ranging from 39.8 to 50.3 g) compared to the more variable differences in grain width and weight. Based on these results we hypothesise that the 5A locus increases grain weight by a primary effect on grain length, which confers the potential for further enhancements by pleiotropic effects on grain width. The grain length effect is genetically controlled and stable across environments, whereas the pleiotropic effect on grain width occurs later in grain development and is more environmentally dependent and variable. The final magnitude of the 5A grain weight increase (ranging from 4.0 to 9.3 %) is thus determined by the extent to which the late stage pleiotropic effect on grain width is manifested and the potential exploited. This could explain why the grain width increase was significantly correlated with the increase in TGW (*r* = 0.98, p = 0.004) whilst grain length was not (*r* = 0.71, p = 0.18).

By dissecting TGW to a more stable yield component (grain length) we were able to classify RILs in a qualitative/binary manner (i.e. “short” or “long” grains) which enabled the fine mapping of the 5A locus to a genetic distance of 4.3 cM. We identified roughly 400 genes in this interval that are expressed in the grain, several of which have annotations associated with genes implicated in the control of grain/seed size. Although it is premature to speculate on potential candidate genes, identification of the causal polymorphism will provide functional insight into the specific mechanism by which pericarp cell size and grain weight are controlled in polyploid wheat.

The consistent effect of the 5A locus on grain length and weight did not always translate into increased yield. In the original DH analysis, the 5A TGW effect co-located with final yield in seven of the twelve environments. This overall positive trend was also reflected in the NILs, although yield increases were only significant in 2014. We concluded that the effects on yield are modulated by a series of smaller negative effects on other yield components which could be due to additional genes within the broader 5A region. Alternatively, it could be that the full potential of the grain length effect will be realised only under certain environments or in combination with other genes.

By understanding the biological mechanism by which the 5A locus achieves increased grain size, hypotheses can be generated to combine genes in an informed and targeted way. For example, we are combining the 5A grain length/pericarp cell expansion effect with the *TaGW2* mutants which affect grain width (presumably through pericarp cell proliferation) to determine if they act in an additive or synergistic manner. Identifying the 5A gene will also allow the function of the homoeologous copies on chromosomes 5B and 5D to be determined. This is important since the effects of grain weight QTL in polyploid wheat are often very subtle compared to those in diploid species (Borrill *et al.,* 2015; Uauy, 2017). Modulating the function of all three homoeologs simultaneously holds the potential to expand the range of phenotypic variation and achieve effects comparable to those in diploids e.g. *NAM-B1* (Uauy *et al.,* 2006; Avni *et al.,* 2014; Liang *et al.,* 2014). Ultimately, identifying the genes and alleles that control specific yield components and understanding how they interact amongst them and with the environment will allow breeders to manipulate and fine-tune wheat yield in novel ways.

## Acknowledgements

This work was supported by the UK Biotechnology and Biological Sciences Research Council (BBSRC) grants BB/J003557/1 and BB/J004588/1. JB was supported by the UK Agriculture and Horticulture Development Board (AHDB) and the John Innes Foundation. We thank Ricardo Ramirez-Gonzalez for the BLAST of iSelect SNPs to the IWGSC 1.0, David Swarbreck and Gemy Kaithakottil (Earlham Institute) for *In slllco* mapping of TGACv1 gene models to the IWGSC RefSeq1.0, the JIC Field Trials Horticultural services for technical support in glasshouse and field experiments.

## Author contributions

JB conducted the developmental time courses, fine-mapping, analysed the data and wrote the manuscript. JSi developed the germplasm used in this study, performed phenotypic assessments and QTL analyses, analysed the data and wrote the manuscript. FM conducted cell size measurements. MLW led the mapping of the Charger x Badger DH population. JSn coordinated and conceived the DH population field trials. CU conceived the experiments, analysed the data and wrote the manuscript. All authors read and approved the final manuscript.

## Supporting information

Table S1: Developmental traits of 5A NILs

Figure S1: Example of SEM image for cell size measurements

Table S2: Significant QTL identified for TGW in the Charger x Badger doubled haploid population

Table S3: Significant QTL identified for yield in the Charger x Badger doubled haploid population

Figure S2: Yield MTME graph

Table S4: Spike yield components 10 representative single ear samples of 5A NILs

Figure S3: Distribution of grain width of NILs from whole plot samples

Figure S4: Grain development time course of 5A NILs (2014 field trials)

Figure S5: Grain development time course of 5A NILs (2016 field trials)

Figure S6: Comparisons of cell length within genotypes between different sized groups of grains

Figure S7: Comparisons of cell length between 5A+ and 5A- NILs in 2016

Figure S8: Comparisons of cell number in 2015 and 2016

File S1: TGACv1 genes in the fine mapped interval and associated expression data

